# Single-molecule biophysics experiments *in silico*: Towards a physical model of a replisome

**DOI:** 10.1101/2021.12.01.470844

**Authors:** Christopher Maffeo, Han-Yi Chou, Aleksei Aksimentiev

## Abstract

The interpretation of single-molecule experiments is frequently aided by computational modeling of biomolecular dynamics. The growth of computing power and ongoing validation of computational models suggest that it soon may be possible to replace some experiments out-right with computational mimics. Here we offer a blueprint for performing single-molecule studies *in silico* using a DNA binding protein as a test bed. We demonstrate how atomistic simulations, typically limited to sub-millisecond durations and zeptoliter volumes, can guide development of a coarse-grained model for use in simulations that mimic experimental assays. We show that, after initially correcting excess attraction between the DNA and protein, qualitative consistency between several experiments and their computational equivalents is achieved, while additionally providing a detailed portrait of the underlying mechanics. Finally the model is used to simulate the trombone loop of a replication fork, a large complex of proteins and DNA.

## Introduction

Single-molecule experiments can characterize physical and biochemical properties of individual biomolecules and biomolecular assemblies to provide key insights into the mechanisms that underlie biological function [58, 8, 37, 65, 43, 19]. However, most experimental techniques that operate on individual biomolecules, including force spectroscopy [8, 37] and fluorescent microscopy [44, 52], can simultaneously probe a very limited number of degrees of freedom. Such limitations can be problematic when it comes to interpreting the results of single-molecule measurements because the functional behavior of a biomolecule may depend on the state of one or more “hidden” degrees of freedom, such as the conformation of a protein [34, 52]. Physics-based computational modeling of single-molecule experiments can greatly aid functional interpretation of the measurements by resolving degrees of freedom that are neither controlled nor observed in the experiment.

Among the most popular computational approaches, explicit solvent all-atom MD simulations have been applied widely to study the molecular mechanisms underlying biological and biotechnological systems [18, 27, 21, 14, 31]. However, the explicit representation of each atom limits routine application of the method to small systems and/or relatively short timescales. Modeling larger systems for longer timescales requires dramatic reduction of the degrees of freedom, or coarse-graining, of a system. In contrast to all-atom simulation, where there are relatively few models available that are all general in nature and have each been developed using similar approaches over decades, coarse-grained (CG) models are extremely diverse in their parametrization and domain of application.

The parametrization of a CG model can, in general, be classified as either top-down or bottom-up. Many CG models have been parameterized using a top-down approach, where the interaction sites and the energy potentials associated with them are chosen using a mixture of intuition, physical law, and trial-and-error until the model produces desired, usually experimentally-measured, properties for some test systems. Examples of such models include: the general-purpose Martini model [7]; the oxDNA [39, 55] and 3SPN2 [13] models for DNA; the AWSEM structured protein model [6]; the various models for disordered peptides from the Hummer [20], Onck [10], Best [63], Papoian [61], and Mittal [48], labs; rigid body and elastic network models of proteins used in virus capsid assembly studies [38, 25, 12]; and rigid body protein models of the bacterial cytoplasm from the Elcock lab [33] and of chromatin from the Schlick lab [2], among others. The Levy lab developed a top-down model with emphasis on electrostatic and aromatic ring interactions to predict the binding geometry of a DNA strand to single-stranded DNA binding protein (SSB) proteins of several species, showing good agreement with available crystallographic structures [36, 40].

At the opposite end of the spectrum for model development, bottom-up coarse-graining starts with simulating a detailed model of the system, typically all-atom, to gather statistics. After selection of CG interaction sites, one of several approaches is used to optimize the CG interaction potentials until agreement of forces, configuration, or general thermodynamic observables is obtained between the CG model and the detailed model. CG potentials are commonly split into bonded and non-bonded parts with non-bonded potentials usually chosen to be pairwise. Collective variable-based potentials, for example those dependent on density, are sometimes used [28, 60]. Pairwise force-matching optimization strategies include the simplex algorithm [35], iterative Boltzmann inversion [42], inverse Monte Carlo [29], and machine learning [11], among others [56].

Compared to top-down, a bottom-up model will precisely capture the effective interactions between CG sites, and arguably it may be less biased by the expectations of the model developer. On the other hand, a bottom-up model may be criticized for inheriting bias from the underlying detailed model and for lacking predictive power because the model primarily reproduces what is already known from simulation of the underlying detailed model. Nevertheless, there are cases where bottom-up coarse-graining has been successfully applied to investigate systems at length scales and timescales much larger and longer than those that could be studied with an underlying all-atom model [33, 12, 59, 57]. For example, our own CG model of ssDNA [32] was parametrized using 10 microseconds of all-atom MD simulation of a 60-nucleotide (nt) DNA strand performed on a supercomputer, yet it enabled workstation-powered studies of a 1000-nt DNA strand transport through a dual nanopore systems [3] for durations of hundreds of microseconds, the simulations presently impossible to carry out using an explicit-solvent atomistic description. However, it should be noted that bottom-up models will inherit any deficiencies of the underlying higher resolution model. Thus, in the case of our ssDNA model, introducing weak non-bonded repulsion was needed to swell the DNA strand to a size consistent with small-angle x-ray scattering experiments [49, 32].

Here we demonstrate the extension of our CG ssDNA model to include SSB, a globular protein that plays a key role during DNA replication and repair processes by preventing secondary structure in the nascently unwound ssDNA and by recruiting repair proteins to sites of DNA damage. We model the protein as a rigid body, reducing the degrees of freedom for each SSB molecule to six. The interactions between CG ssDNA beads and the SSB were obtained using the iterative Boltzmann inversion (IBI) method to match structural distributions from all-atom simulations. However, we found that the binding free energy of a short DNA fragment was overestimated, and so we tuned the interaction potential until the model reproduced the experimental binding free energy. Thereafter, quantitative agreement with the experimentally measured elasticity of a DNA–SSB complex emerged without further refinement. The model was then used to study competition between DNA fragments and SSB molecules in systems that mimic experimental assays. Finally, we introduced SSB– SSB interactions using a top-down approach and used the model to investigate the possible structure of a trombone loop within the DNA replication fork.

## Results

Seeking access to timescales that would enable *in silico* modeling of constructs used in experiments to probe SSB–ssDNA and SSB–SSB interactions, we set out to construct a coarse-grained model of SSB using all-atom MD simulations to guide the process. The construction of the model is described below.

### Beads-and-grids models of ssDNA-SSB assembly

To model an ssDNA–SSB complex, we begin by selecting an existing CG model of ss-DNA that reproduces the experimentally-measured elastic properties of DNA, namely a model developed several years ago by our lab [32]. In that model, each nucleotide is represented by two beads. The interaction potentials were parametrized iteratively to match target structural distributions obtained from all-atom simulations, including bond and angle distributions and pair distance distributions. A small correction to the resulting non-bonded potentials was applied to swell the CG DNA so that its radius of gyration matched that found in experiment, presumably correcting a systematic bias in the underlying all-atom force field. The final version of the model quantitatively reproduced the elastic response of a DNA molecule to applied tension [51, 47], although it was not specifically parameterized to do so.

The successful application of the iterative coarse-graining procedure for constructing our ssDNA model led us to apply the same approach to SSB. First, we chose to represent the protein as a rigid body without internal degrees of freedom because conformational fluctuations of the core SSB protein were seen to be small in all-atom simulations [30]. To support the iterative Boltzmann inversion procedure, we decided to use tabulated three-dimensional grid-based potentials to describe the interactions between each CG DNA bead type and the protein. Figure 1 provides a pictorial representation of the all-atom and CG model of an ssDNA–SSB complex. Note that, in building our model of SSB, we have neglected its intrinsically disordered C-terminal tails which play a role in recruitment of auxiliary replication and repair proteins [1, 53, 23, 22]. While adding explicit representation of the flexible tails to our model is straightforward from a technical standpoint, the parameterization of the model, in particular, of the tails’ interactions with auxiliary proteins and with other tails, is challenging because of the lack of the required experimental data.

**Figure 1:**
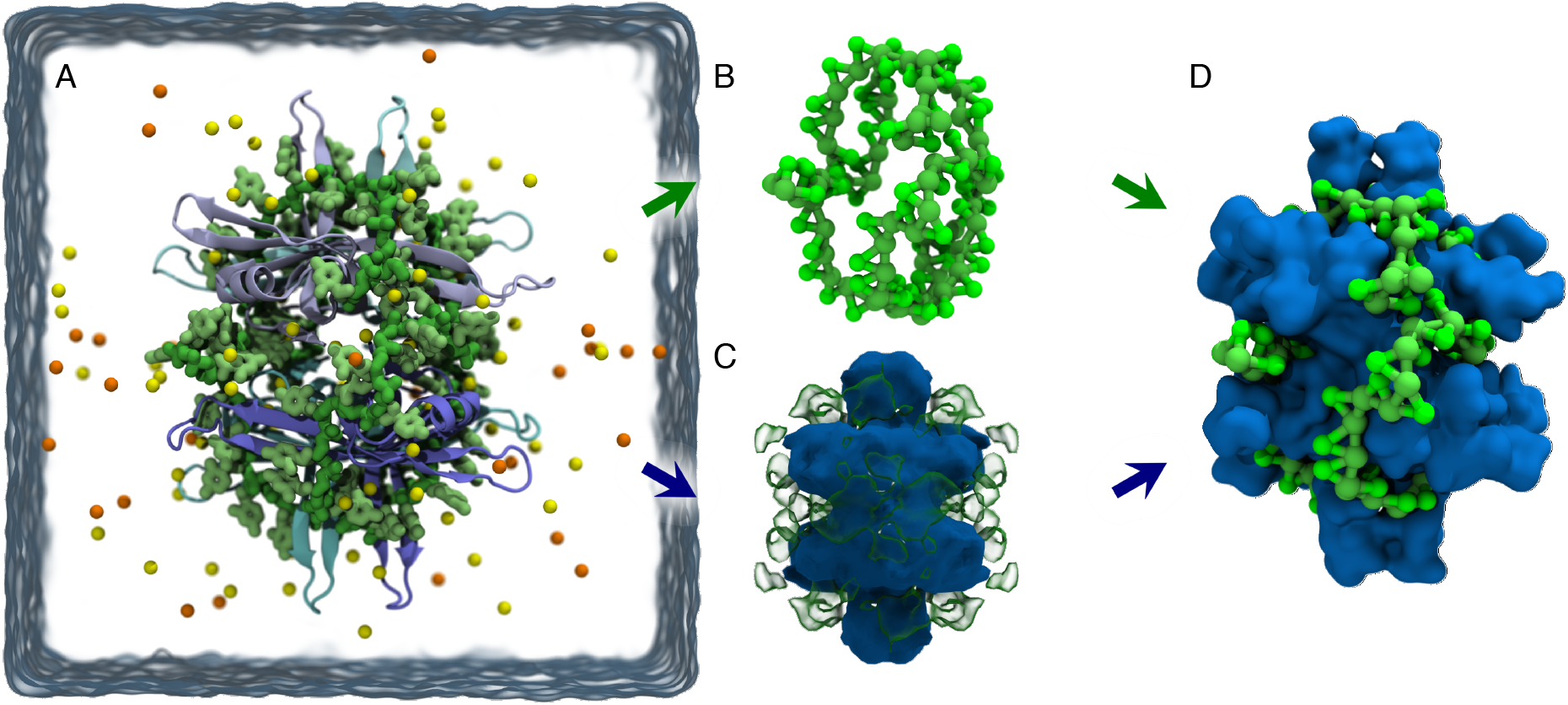
Beads-and-grids model of ssDNA-SSB assembly. (A) All-atom model of SSB (blue and cyan) bound to ssDNA (green) in a 120 mM electrolyte solution (semi-transparent surface) containing cations (Na^+^; yellow) and anions (Cl^-^; orange). (B) Beads-on-a-string representation of the ssDNA molecule depicted in panel A. Two beads, P (phosphate; green) and B (base; light green), represent each nucleotide. The beads interact through a set of tabulated potentials that were optimized to reproduce distributions extracted from atomistic simulations and the experimentally inferred radius of gyration of unstructured ssDNA. Solvent is represented implicitly through the non-bonded bead-bead interaction potentials. See Ref. 32 for additional details. (C) Grids-based model of an SSB particle. The physical SSB particle features a mass and a moment of inertia and can translate and rotate in response to external forces and torques. For each type of the CG beads used in the ssDNA model, a three-dimensional potential grid is tethered to the SSB particle to prescribe the interaction between the protein and that bead type. A cross-section of the potential for P beads is depicted. At a given timestep, for each force applied to an ssDNA bead, the opposite force and corresponding torque are applied to the SSB particle. (D) Beads-and-grids representation of the SSB–ssDNA complex.

### Refinement of the model against all-atom simulation data

To generate target distributions required for the parametrization of our CG model, we probed the interactions between SSB and short fragments of ssDNA using the all-atom MD method. A typical simulation system contained 31 fragments of dT_5_ distributed around an SSB molecule in a cubic volume of 100 mM NaCl electrolyte, Figure 2A. Five replicate systems were constructed differing by the initial placement of the fragments and run in parallel, generating trajectories that amounted to 14 *μ*s of aggregate simulation time. During each simulation, the protein was restrained to its initial coordinates while the DNA fragments were observed to spontaneously bind to and dissociate from the SSB surface, Supplementary Animation 1. The resulting simulation trajectories were then mapped into CG DNA trajectories as described in Ref. 32, Figure 2B. From the CG representation of the all-atom trajectories, the three-dimensional, position-dependent target density was computed for each type of the DNA beads and averaged over the symmetry axes of the SSB homotetramer, Figure 2C.

**Figure 2:**
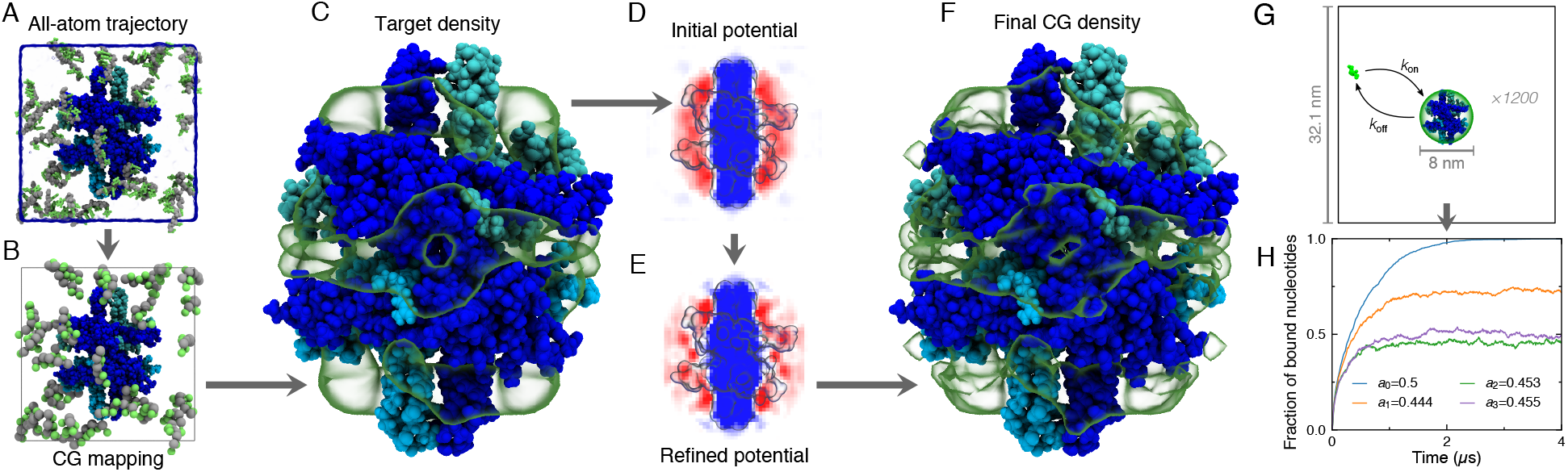
Parametrization of the CG model of SSB. (A) Generation of initial target data for parametrization through all-atom MD simulations of SSB surrounded by 31 dT_5_ ssDNA fragments. The SSB protein was restrained to its initial coordinates. (B) Atomic coordinates of the ssDNA fragments from the MD trajectories were mapped into a CG representation. (C) Target density of each CG ssDNA bead type was extracted from the mapped trajectory. (D-F) Iterative Boltzmann inversion of the target density. Boltzmann inversion provided an initial estimate for the CG SSB–ssDNA interaction potential (D). Each CG potential was refined (E) until they produced a CG density (F) that matched the target density. The potentials and densities shown are for the P ssDNA beads. (G) Refinement of the CG SSB model against experimental binding affinity. At a given moment, a DNA fragment was considered bound if any of its beads were inside the green sphere surrounding the SSB. In the *i*th step of refinement, 1,200 independent simulations were performed of a dT_8_ fragment interacting with the SSB potentials with the negative portion of the SSB potentials scaled by *α*_*i*-1_, see text for a detailed description. (H) The fraction of DNA nucleotides bound to SSB as a function of simulation time. The *α_i_* value indicate the scaling factor applied in the simulations after the *i*th refinement of the interaction potential. According to experiment, the fraction of bound nucleotides in the simulated system should be 0.5.

We began parameterization of our CG model of the ssDNA–SSB assembly by building six CG models of the above all-atom systems matching their nucleic acid composition and dimensions. The SSB protein was represented using a set of trial grid-based potentials that acted on the ssDNA beads; the initial guess for the potentials was obtained through Boltzmann inversion of the target densities, Figure 2D. The trial potentials were applied to the CG DNA beads during subsequent simulations performed using the ARBD software package [4] developed in-house. The resulting CG trajectories, lasting a total of ~ 180 ns, were analyzed to produce density maps of ssDNA CG beads. For a given bead type, the ratio of the CG density to the target density provided a space-dependent correction to the SSB potential

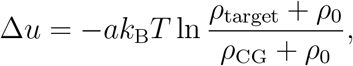

where the scaling factor *a* = 0.1 ensured gradual convergence and *ρ*_0_ = 10^-6^ beads/Å^3^ protected against numerical instability. Note that one nanosecond of a CG simulation corresponds to considerably longer real-world time interval because of the smoothing of the interaction potentials [66]. By repeating the process for a total of 225 iterations, CG potentials (Figure 2E) were obtained that produced CG densities (Figure 2F and Supplementary Animation 2) in excellent agreement with the target densities.

### Refinement of the model against experimental data

Experiment determined the binding affinity between SSB and a dT_8_ molecule to be *K*_eq_ = 20, 000 M^-1^ [24]. This binding affinity corresponds to a 50% occupancy of the bound state for a single pair of ssDNA and SSB molecules confined to a cubic volume 32 nm on side. To test and refine our model against the experimental binding affinity data, we constructed 1,200 CG systems each containing one fixed SSB particle at the center of a cubic volume (32 nm on side) and one dT_8_ ssDNA fragment, Figure 2G. The systems were simulated in parallel for 4 *μ*s each.

In the simulations carried out using the SSB potentials derived by matching the all-atom densities, the fraction of bound nucleotides increased until nearly all DNA fragments became bound to the SSB, Figure 2H. This behavior indicates excessive attraction of ssDNA to the SSB, which we interpret as a manifestation of the artifact in the all-atom MD force field that we used to derive the target ssDNA density.

To bring the affinity between ssDNA and SSB in agreement with experiment, we scaled the negative values of the SSB particle potential for the P beads of ssDNA with a scaling factor *α*_1_ = [1 — (*f*_0_ – 0.5)], where *f*_0_ is the time-averaged fraction of bound nucleotides. We chose to scale only the negative values of the potential because the positive values describe the effect of steric repulsion. The simulations of dT_8_ binding were repeated using the new SSB potentials to obtain a new estimate for the binding fraction, *f*_1_. This process was repeated iteratively with the scaling factor adjusted as *α_i_* = *α*_*i*-1_[1 – (*f*_*i*-1_ – 0.5)]. After only three additional iterations, the fraction of DNA nucleotides bound to the SSB was well converged to the experimental value of 0.5, see Figure 2H.

Upon completion of the above refinement procedure, the potentials reproduced the experimentally determined binding fraction of dT_8_ nucleotides, and therefore the standard binding free energy. In subsequent simulations, we modeled the SSB protein in ARBD as a rigid body having a mass and moment of inertia determined from the all-atom model. At each timestep, the force vector applied to each ssDNA bead by the grid was multiplied by –1 to obtain the force 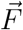 applied by the bead on the SSB. The corresponding torque 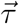 was obtained from 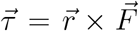, where 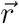 is the vector from the center of the SSB to the position of the point particle. The force and torque on the SSB from all DNA beads were accumulated, allowing integration of the equations of motion for both translational and orientational degrees of freedom [9].

### Experimental validation of the model

Force spectroscopy, usually employing one or more optical or magnetic traps, can be used to probe the mechanical properties of a biomolecular complex. The mechanical work required to disassemble a binary complex of biomolecules, which is equivalent to the binding free energy of the complex, can sometimes be inferred from experimentally measured force-extension curves. The Chemla lab characterized the free energy landscape of SSB binding to ssDNA using an experimental assay depicted schematically in Figure 3A,B [54]. The extension of a DNA construct under tension is measured before (X_ssDNA_) and after (X_SSB_) SSB binding. The difference between these two extensions, X_ssDNA_ – X_SSB_, indicates how much the DNA construct shrinks due to SSB binding. At low tensions, both X_ssDNA_ and X_SSB_ approach zero, and the difference between them is small. Under a moderate tension (~ 5 pN), the DNA is stretched against an entropic force. The DNA molecule exhibits less extension when SSB is bound to it because the ssDNA bound to the SSB is not unavailable for stretching. At slightly higher forces, SSB partially unbinds from ssDNA. At even higher forces, SSB dissociates from the DNA and X_ssDNA_ – X_SSB_ approaches zero.

**Figure 3:**
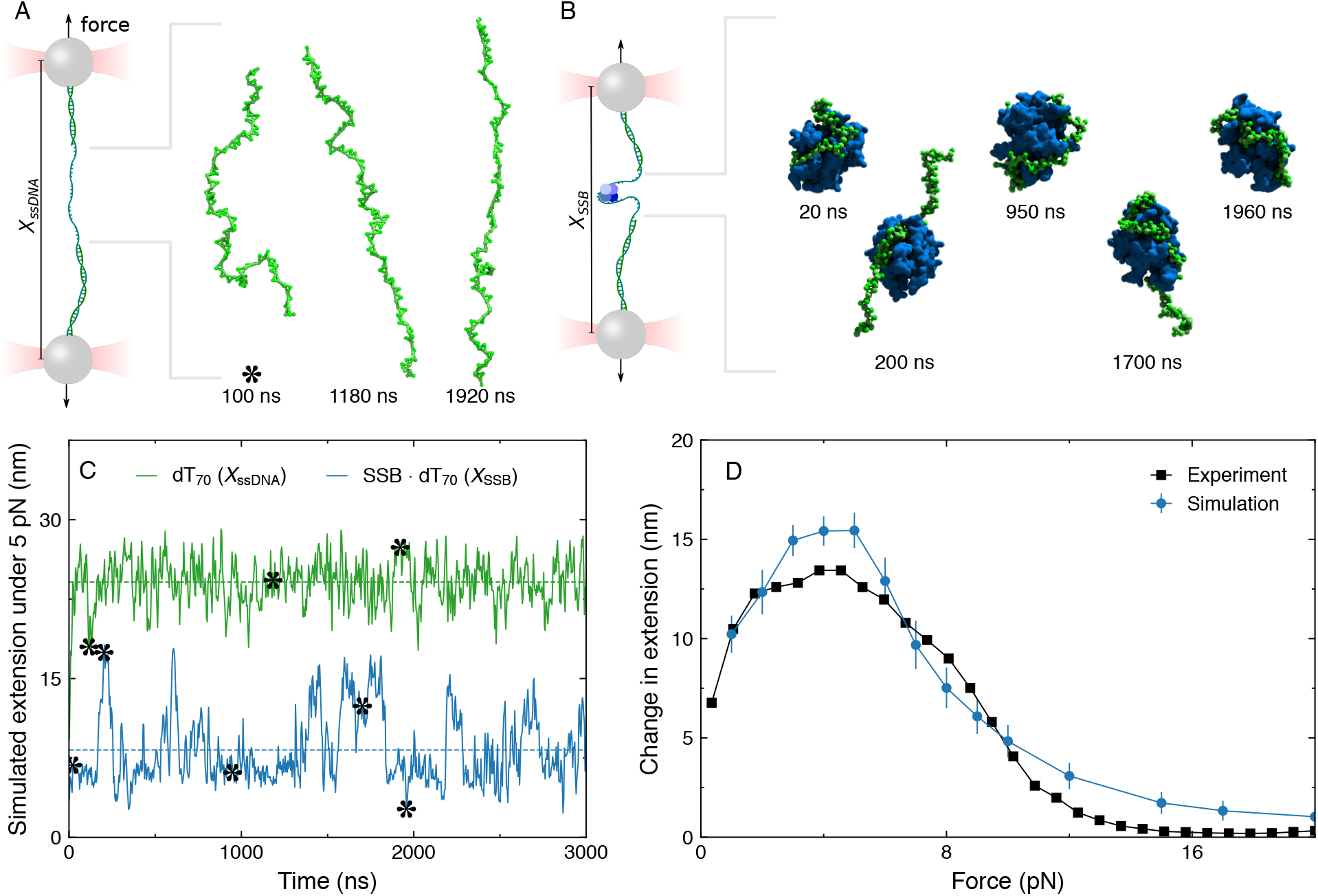
Force-extension dependence of an ssDNA–SSB complex measured *in vitro* and *in silico.* (A) Schematic of a single-molecule experiment including a pair of optical traps that has been used to study SSB [54]. A 5 pN force was applied in opposite directions to the ends of a 70-nt ssDNA molecule in CG simulations representing the experiment. Exemplary configurations of the DNA are depicted on the right with the DNA represented by small green spheres connected by rods. (B) Simulation trajectory mimicking the experiment. A 5 pN force was applied in opposite directions to the ends of a 70-nt ssDNA molecule while an SSB protein was bound to the ssDNA. The extension of the ssDNA increases and decreases as the SSB molecule wraps and releases the ssDNA. Exemplary configurations of the ssDNA–SSB complex are depicted on the right, with the protein represented as a blue molecular surface. (C) Simulated extension of ssDNA in complex with SSB (blue) and in isolation (green). The asterisks denote the moments of the trajectory depicted in panels A and B. The dotted lines indicate the corresponding trajectory average values. (D) Average change in dT_70_ extension upon SSB dissociation experimentally measured in *vitro* (black) and in *silico* (blue) as a function of applied tension. Each simulated data point was obtained by time-averaging DNA extension and taking the difference between the values obtained with and without SSB. Experimental data were provided by the Chemla lab [54]

To test how well our CG model can reproduce the results of single-molecule experiments, we performed force-extension experiments on the SSB–ssDNA complex *in silico*. In each simulation, a dT_70_ molecule was stretched by applying opposite constant forces to the P beads at each end of the molecule. First, the simulations were performed in the absence of SSB, Supplementary Animation 3. The extension during these simulations reached a steady state value within ~ 50 ns. Low, average, and high extension configurations under a 5 pN force are shown in Figure 3A and a typical extension trace is plotted in Figure 3C.

We began our simulations of the force-extension dependence of an ssDNA–SSB complex from a configuration where ssDNA was wrapped around the SSB, Supplementary Animation 4. Complete dissociation of SSB from ssDNA was not observed, even at the highest applied tension studied (20 pN). Within 50 ns of the beginning of each simulation, the ssDNA extension reached a steady state. The protein–DNA complex was seen to be highly dynamic in the simulation, in particular when the tension force was near 5 pN. Low, average, and high extension configurations under a 5 pN tension are shown in Figure 3B along with an extension trace, Figure 3C.

For each simulation, the extension of the ssDNA molecule was obtained directly from the trajectory coordinates and then time-averaged to yield a single value. The difference between extensions without and with SSB as a function of applied force yielded a curve that could be directly compared to the experimentally obtained curve, Figure 3D. Although our model was not parameterized to reproduce the behavior of an SSB–ssDNA complex under applied force, we observed very close quantitative agreement between our simulation results and experiment.

### Kinetics of SSB-ssDNA interactions

The process of coarse-graining effectively smoothens the interaction potentials, resulting in faster kinetics [66]. For the lengths of ssDNA considered in this study, the rate of end-to-end collisions of an ssDNA molecule is 5 to 50 times higher than in the corresponding experiment [32], whereas the time scale of free diffusion of an SSB particle is solution is exactly same as in experiment, prescribed by the diffusion constant. However, the dissociation rate of an 8-nucleotide ssDNA fragment from the SSB particle was found to be 3000 times greater in our CG simulations than measured experimentally [24]. The difference in the rates of relative speed-up observed for different components of the CG system may impact the quality of the kinetic data extracted from the simulations. The thermodynamic quantities, however, are not expected to be affected by the time scale differences.

Besides resulting in a large boost in the rate of SSB diffusion along the ssDNA molecule, the artificially accelerated dynamics at the SSB–ssDNA interface causes rapid transitions between wrapping geometries of ssDNA on the SSB surface, Figure 4A,B. The extension of SSB-bound ssDNA under 5 pN tension was seen to change in discrete steps in single-molecule experiments [54], while, in our simulations, the transitions occurred too quickly for the steps to be clearly resolved, Figure 4B. Nevertheless, the likelihood of the protein occupying a given wrapping geometry is a thermodynamic property that is expected to be accurately captured by the model. Under tension, the most prevalent wrapping geometry at low force includes ssDNA making a “C” shape on the SSB surface, in contrast with the baseball-seam wrapping geometry suggested by the x-ray crystallographic structure [41]. As the force increases, it becomes increasingly likely that the SSB loses contact with one or more of the four high-affinity binding sites, also called oligonucleotide binding (OB) folds, Figure 4C.

**Figure 4:**
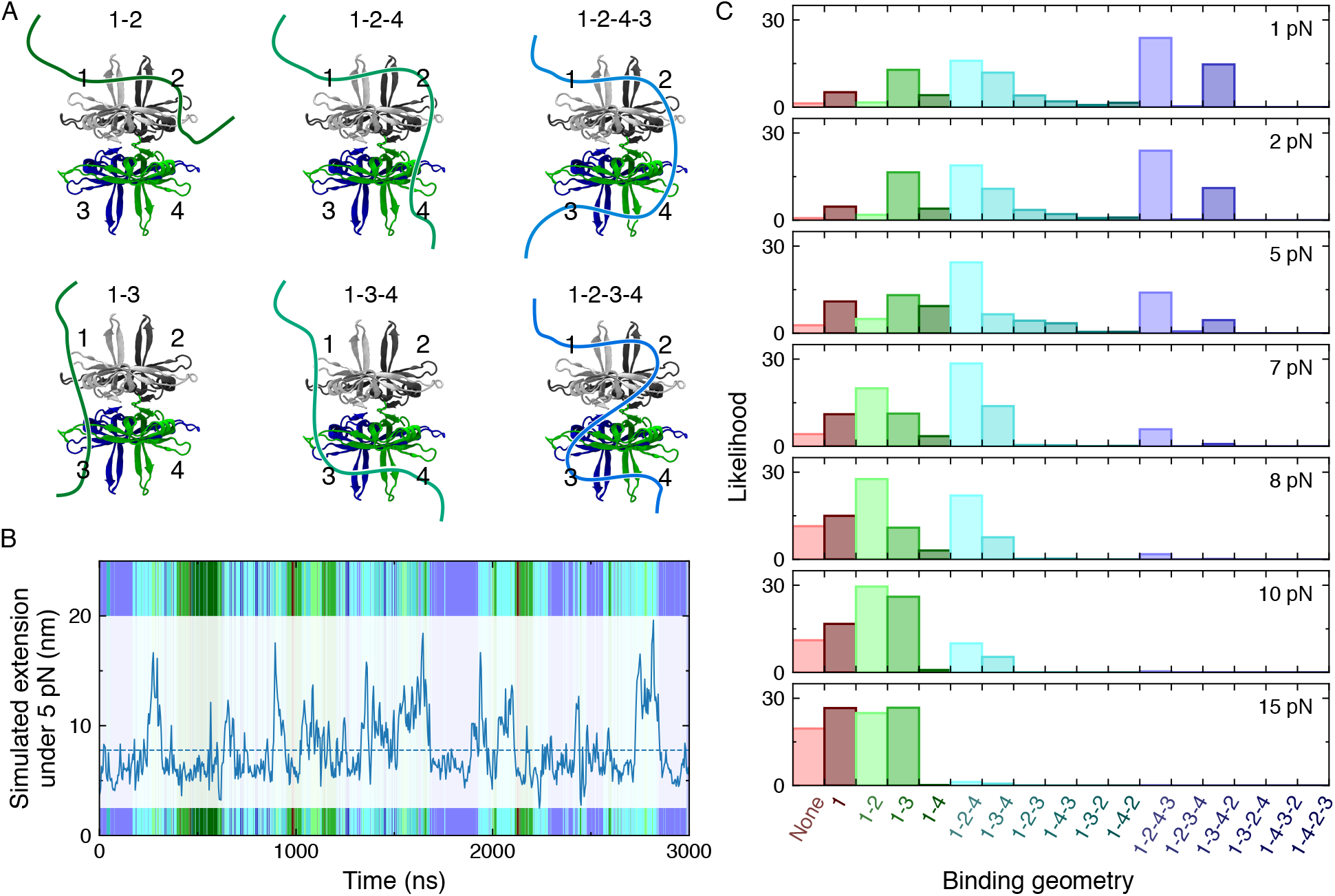
Wrapping geometry of ssDNA on SSB observed in CG simulations. (A) Labeling scheme used to classify the wrapping geometry. Each of the four high affinity OB folds is assigned an integer. The wrapping geometry can be specified by the sequence in which the DNA strand contacts the OB-folds. The symmetry of the SSB tetramer allows the first OB fold contacted by the 5’ end of the strand to be labeled “1”. (B) Wrapping geometry during simulation of SSB–dT_70_ complex under 5 pN tension. The blue trace depicts the instantaneous extension of the complex. At each moment in time, the color in the background of the plot indicates the wrapping geometry using the same color scheme specified in panel C. (C) Force-dependence of the wrapping geometry. The likelihood of the system occupying a given wrapping geometry is shown for the tension force ranging from 1 to 15 pN. At low force, wrapping geometries where ssDNA contacts all four OB folds are most likely. At high forces, wrapping geometries with only two OB fold contacts are more likely.

### Transfer of SSB from one DNA strand to another

Single-molecule experiments can be designed to probe biologically important behavior of biomolecules. For example, DNA replication enzymes generally operate only in the 5’-to-3’ direction so that replication on one of the strands is performed in short (~ 500 bp) sections between priming sites called Okazaki fragments. The ssDNA exposed in these fragments is coated by SSB proteins that bind tightly, yet almost paradoxically must easily be repositioned as the polymerase causes the ssDNA loop to shrink. Analysis of the diffusion of a fluorescently labeled SSB molecule demonstrates that the motion of SSB bound to a long ssDNA construct can be so fast that SSB must be hopping along different, non-continuous fragments of the same ssDNA construct [26], and not sliding or rolling [45, 64]. Subsequent single-molecule experiments characterized the kinetics of exchanging one DNA strand bound to an SSB molecule for another [62]. These experiments demonstrate that SSB can jump from one region of ssDNA to another within the replication fork, allowing it to move out of the way of an active polymerase. The experiments primarily tell us that the exchange process happens, but it remains difficult to determine the steps that typically occur during the exchange, including such details as the structure of the encounter complex where two strands simultaneously bind one SSB protein.

Our CG model of SSB and ssDNA enables simulations of such spontaneous transfer processes, which are much too slow to simulate using the all-atom MD approach. As a demonstration, we constructed a system containing a complex of SSB and dT_50_ and a free dT_40_ molecule, confining the free DNA molecule to reside within a sphere of a 25 nm radius using a half harmonic potential (spring constant *k* =1 kcal mol^-1^ Å^-2^ per bead), whereas the center of the SSB was confined to a smaller, 10 nm radius sphere (*k* = 8 kcal mol^-1^ A^-2^), Figure 5A. The incumbent DNA strand was not confined and was free to leave the immediate volume around the SSB, effectively preventing rebinding. The confinement potential creates a high effective concentration of ~ 25 *μ*M for the invading dT_40_ molecule, which enhances the rate at which an encounter complexes form. During the simulation, we observed the 40-nt strand to bind to the SSB and compete for high-affinity area at the SSB surface until either the invading strand dissociates, Figure 5B, or the incumbent strand dissociates Figure 5C. The strand exchange process is well characterized by the timeseries of the number of nucleotides of each strand bound to the SSB, Figure 5D.

**Figure 5:**
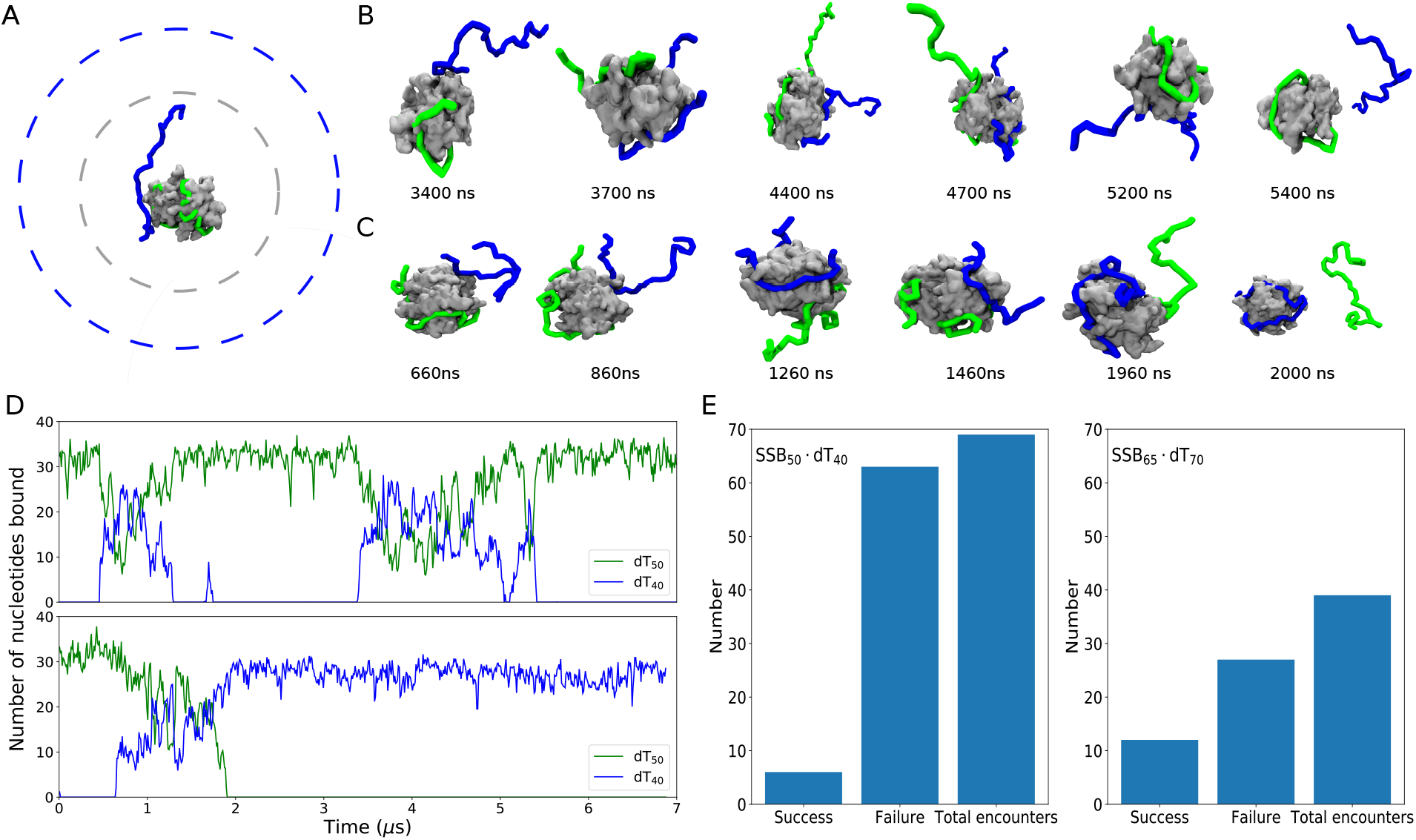
Transfer of SSB between two different-length DNA strands. (A) System containing SSB and two DNA strands, one initially in complex with SSB (green; 50 nt) and one invading strand (blue; 40 nt). The SSB is shown as a gray molecular surface, and the dashed lines represent spherical potentials used to confine the SSB (gray) and DNA strands (blue). (B) Encounter that results in the dissociation of the invading strand. (C) Encounter that results in the dissociation of the incumbent strand. (D) Number of nucleotides bound to SSB from each strand during the simulations depicted in panels C and D. The number for initially-bound and invading strands are respectively shown using green and blue lines. A nucleotide was considered bound to the SSB if a P bead of ssDNA was found within 1-nm of the non-hydrogen atoms of SSB. The CG representation of SSB was replaced by its all-atom representation for this analysis. (E) Statistical outcome of encounter. By repeating the encounter simulation multiple times, the likelihood of an encounter resulting in a successful transfer of the SSB to the invading strand could be determined. The outcome of an encounter was also determined for dT_65_ invading an SSB–dT_70_ complex.

The process of SSB transfer took less than a day to simulate on a single workstation. Hence, it was feasible to replicate the simulation many times in parallel to generate results that could be expressed in statistical terms. One hundred trajectories were produced, and the timeseries of the number of nucleotides bound to SSB was extracted for each strand in each trajectory. The outcome of attempted transfer events were identified algorithmically as successful or failed, depending on whether the incumbent or the invading strand dissociated from the complex, Figure 5E. Representative simulation trajectories of failed and successful transfer events are highlighted in Supplementary Animations 5&6. Similar simulations were carried out to examine the competition between longer dT_70_ and dT_65_ strands.

The results of our simulations show that the likelihood of forming an encounter complex is lower for longer DNA strands, likely because the high-affinity regions of the SSB surface a better protected by the incumbent strand. Once an encounter complex is formed, it appears that the likelihood of successfully transferring the SSB is sensitive to the difference in nucleotides between the two strands, with dT_65_ displacing dT_70_ ~ 30% of the time and dT_40_ displacing dT_50_ just under 10% of the time. The probability of a successful (*ρ*_S_) or failed (*ρ*_F_) transfer is proportional to the Boltzmann-weighted free energy *U* of all molecular complexes present after one strand dissociates, such that *ρ*_S_/*ρ*_F_ = *e*^*β*(*U*_F_-*U*_S_)^. Assuming that the difference in the resulting free energies can be attributed to the free energy of the extra nucleotides of the binding the SSB surface, we find an energy difference of 1.2 and 2.3 *k*_B_*T*(about 0.24 *k*_B_*T*/nt) for systems with 65 nt and 40 nt invading strands, respectively. The free energy is within the range of values suggested by force spectroscopy experiments [54] that indicate ~ 0.44 *k*_B_*T*/nt for the first 35 nucleotides that bind and 0.07 *k*_B_*T*/nt for the last 9 nucleotides between states corresponding to 56 and 65 nt bound. The total number of direct contacts between ssDNA and SSB was observed to increase from ~ 32 nt for the single dT_50_ to 41 nt upon binding of the invading dT_40_ strand, and from ~ 36 nt for dT_70_ to 42 nt for the invading dT_65_ strand, suggesting that at most 42 nucleotides can directly contact the protein at any given time, regardless of the length and the number of bound ssDNA fragments.

### Exchange of SSB proteins bound to ssDNA

Single-molecule experiments from the Ha lab demonstrated the ease with which SSB can be repositioned or replaced by another protein [46, 62]. In a typical experimental assay, a fluorescently-labeled SSB molecule is initially bound to a surface-tethered DNA construct that had an exposed dT_70_ strand. The construct has its 5’ end of the ssDNA stem from a dsDNA duplex which in turn is tethered to a glass slide, leaving the 3’ end of dT_70_ exposed to solution. When a second SSB molecule encounters the SSB-ssDNA complex, it can replace the incumbent SSB [62]. Three states could be detected via the measurement of the FRET efficiency, corresponding to a single labeled SSB molecule bound to the DNA, two SSBs bound to the DNA, and, finally, with having the unlabeled SSB from the solution bound to the DNA. However, the experiments are unable to detect intermediates that likely occur during the exchange process where two SSBs are simultaneously bound to a single DNA construct.

We recreated this experimental assay *in silico*. In a typical simulations, a DNA construct was tethered to a half harmonic repulsive surface. One SSB molecule was partially wrapped with the ssDNA part of the construct and a second SSB particle was placed 8 Å away from the nearest DNA bead, Figure 6A. To describe the SSB-SSB interactions, we used a previously developed protocol [50] that clusters atoms according to the CHARMM36 [16] Lennard-Jones (LJ) parameters, constructs approximate 3D LJ potential maps, an electrostatic potential map from APBS [15, 17], and maps with the distribution of corresponding LJ and electrostatic charges. Note that the sum over an LJ charge map equals the number of atoms of the corresponding LJ type in the SSB protein; negative LJ charges are never employed. If a voxel of the charge map (generalized, not necessarily electrostatic) of one SSB overlaps with the potential map of the other, a force is computed as the charge of that voxel multiplied by a negative gradient of the potential at that voxel, with the latter being determined using linear interpolation of the potential map. The corresponding torque is computed as a cross product between the vector connecting the center of the SSB (with the charge map) with the center of the charge voxel and the force on the charge voxel. For efficiency, grid-grid interactions were only computed every 20 steps.

**Figure 6:**
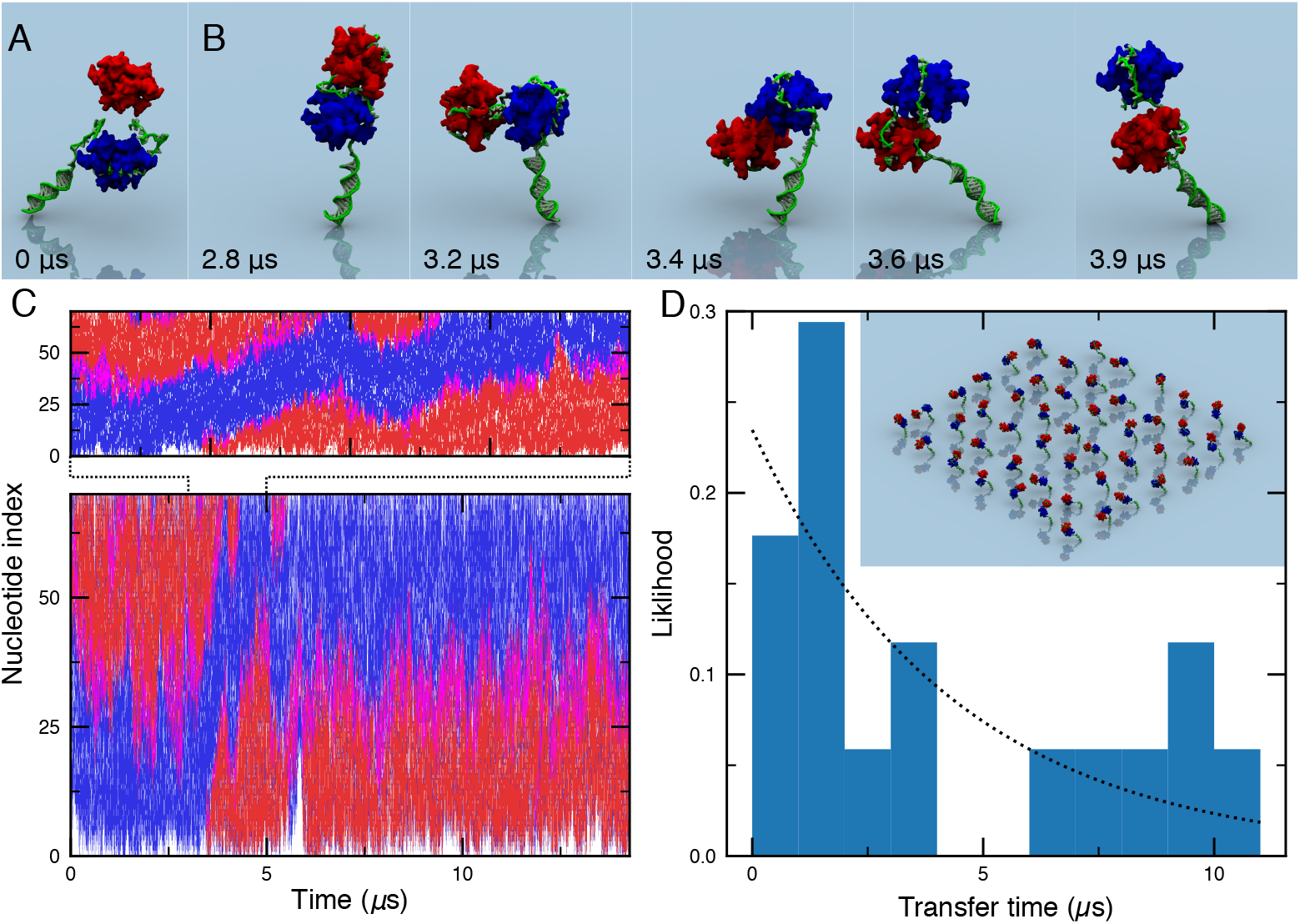
Exchange of two SSB proteins bound to the same ssDNA fragment. (A) System containing a DNA construct tethered to a plane by a harmonic potential with two SSB molecules in close proximity. (B) Snapshots illustrating the simulated process of SSB exchange. (C) Exemplary trace depicting the instantaneous contact status of each ssDNA nucleotide of the DNA construct. For a given configuration, each nucleotide is either bound to the SSB initially placed near the ssDNA–dsDNA junction (blue), to the SSB initially bound to the free 3’ end of the ssDNA (red), to both SSB proteins (magenta), or neither SSB (white). A nucleotide was considered to be in contact with the SSB if its P bead was within 1 nm of any non-hydrogen SSB atoms. The CG representation of SSB was replaced by its all-atom equivalent for this analysis. The top graph shows a zoomed-in view of the bottom graph. (D) Probability of observing an exchange event versus simulation time. The exchange events were detected algorithmically by determining the median index of nucleotides bound to each SSB protein. Identical initial configurations were used to initiate 48 independent simulations, differing by the random generator seed. Multiple transfers events within a continuous simulation trajectory were considered as independent transfer events. The inset illustrates instantaneous configurations of the 48 simulation systems; each system was simulated in isolation from the others. Dotted line shows an exponential fit to the distribution.

Immediately after the simulation started, the second, upper SSB molecule became bound to the ssDNA. After 2.8 *μ*s, some of the bound DNA peels away from the upper SSB molecule (the one initially bound near the free end of the ssDNA), exposing its high affinity DNA binding sites. Around the same time, the flexible DNA construct experiences a thermal fluctuation that positions the upper SSB near to the ssDNA-dsDNA junction, first three panels of Figure 6B. The ssDNA nucleotides adjacent to the junction then spontaneously bind the upper SSB, fourth panel of Figure 6B. For the next three hundred nanoseconds the upper SSB makes contacts with both the free end and the junction end of the ssDNA, third and fourth panel of Figure 6B. Meanwhile, the lower SSB gradually moves towards the free end until finally the exchange is complete, final panel of Figure 6B. The entire exchange process is highlighted in Supplementary Animation 7. Examination of the contacts formed between the ssDNA and the SSBs allows the exchange to be detected algorithmically, Figure 6C. Forty-seven additional replicas of the system, starting from the same initial conformation, were then independently simulated for a similar duration to determine whether the exchange event could be observed repeatedly and to obtain an estimate the exchange rate.

In each simulation, both SSBs were constantly in contact with the ssDNA; complete dissociation of an SSB molecule was not observed, though multiple exchanges were observed in individual simulations. Taken together, the simulations provided a statistical view of the exchange process within the intermediate complex, allowing the rate of exchange to be determined (~ 0.23 *μ*s^-1^) from a fit to the dwell time distribution, Figure 6D. We stress, however, that the absolute exchange rate determined from our CG simulations should be interpreted cautiously because of the smoothing of the underlying free energy landscape, which results in a process-dependent acceleration of the system’s dynamics [66]. Rather than examining the absolute rates, such simulations could be used to compare the exchange rates, for example, probing how the exchange rate depends on the length of the ssDNA or perhaps on modifications of the SSB structure. Our simulation approach could be generalized to model competitive binding of SSB and other DNA-binding proteins, such as RecA, to the same DNA fragment, providing insights into the microscopic mechanisms of DNA repair [31].

### Model of the T7 bacteriophage trombone loop saturated with bacterial SSB

According to the current model of DNA replication in prokaryotes and some viruses, the parental dsDNA is unwound by a helicase that is in complex with a pair of DNA polymerase molecules. One polymerase replicates the leading strand, traversing the DNA in the same direction that the helicase moves, while the other polymerase replicates the lagging strand in the opposite direction, Figure 7A. The lagging strand polymerase therefore must “back-stitch”, replicating the DNA in a piecewise manner while the “trombone” loop grows, on one end, from ssDNA released by the helicase and, on the other, from daughter dsDNA at the polymerase. The main biological function of SSB proteins is to protect the lagging strand exposed during DNA replication from degradation by endonucleases and secondary structure formation.

**Figure 7:**
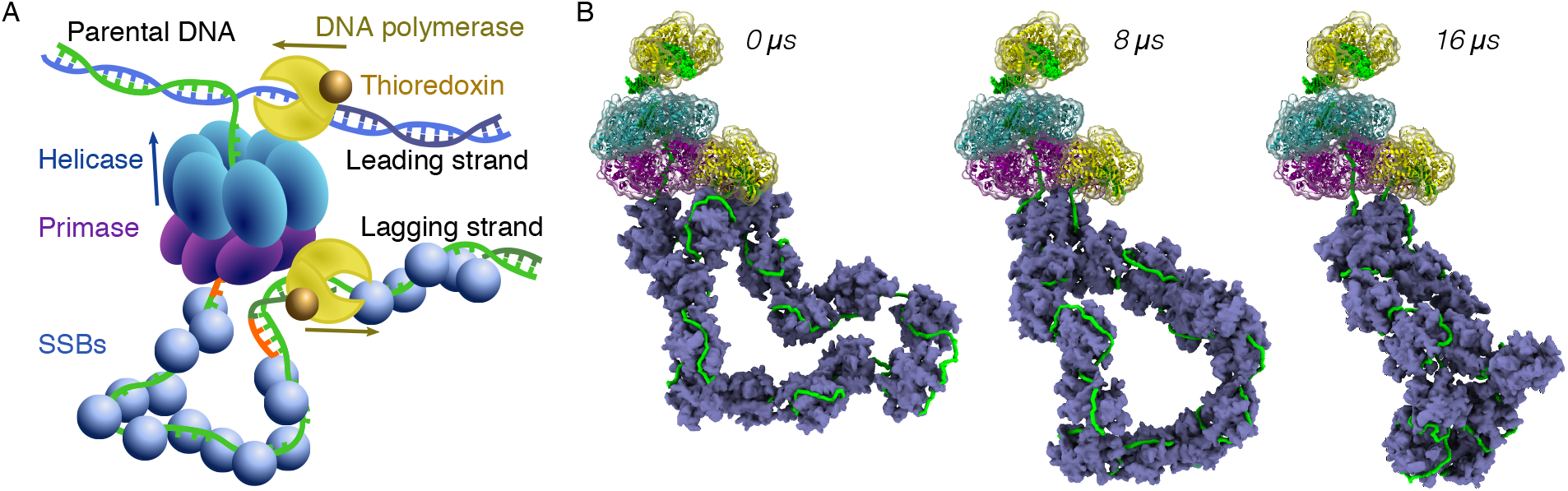
Physical model of the trombone loop of the DNA replication fork. (A) Schematic depicting the T7 DNA replication fork. (B) Snapshots of the simulation of the trombone loop. Semi-transparent molecular surfaces represent the helicase (teal) in complex with the primase (purple) and polymerase (yellow) proteins, which could be resolved in cryo-EM and are represented in the simulation by grid-based potentials that act on the ssDNA and SSB proteins. Double-stranded DNA and ssDNA beads bound to the above proteins are depicted but were not represented explicitly in the simulations. The remaining ssDNA (green beads and rods) is coated with SSB proteins, each drawn as a purple molecular surface.

We constructed a physical model of SSB proteins in the context of the replication fork, using the minimalist replication machinery of the T7 bacteriophage as our target system. We manually docked structured parts of the helicase, primase, thioredoxin and polymerase proteins into the cryo-EM map of the replication fork and used the procedure described in the previous section for SSB–SSB interactions to convert the atomic models into potential maps that would act on the SSB rigid body proteins. In addition, we used the cryo-EM map as a potential to sterically repel ssDNA from the structured parts of the replication fork. We then constructed a model using these potential maps with a 500-nt loop of ssDNA saturated with 14 SSB molecules to represent the trombone loop of the replication fork. The model of the loop was constructed by applying transformations to a previously equilibrated model of the ssDNA-SSB construct. The DNA loop was anchored at locations resolved by cryo-EM near the primase on the one end and the lagging-strand polymerase at the other. Our model represents the state of the replication fork immediately after the lagging polymerase binds a primed template strand and as such does not include an explicit bead-based representation of the dsDNA. To the best of our knowledge, this is the first structurally correct physical model of a trombone loop.

During the 16 *μ*s simulation, all SSB proteins remained bound to the ssDNA of the trombone loop, whereas the loop as a whole was seen to undergo significant rearrangements, Figure 7B and Supplementary Animation 8. Most prominently, the loop became more compact with SSB proteins frequently making contact with more than one region of the ssDNA strand, effectively sharing the strand. Interestingly, a single SSB protein became stably bound to both ends of the loop where it enters the structured region of the replication fork, giving a straightforward prediction to be tested experimentally.

## Discussion

Using a bottom-up approach, we have developed a CG model of a structured DNA-binding protein, SSB, compatible with a previously developed 2-bead-per-nucleotide model of ssDNA. The key methodological innovation of our approach is that the protein is modeled as a rigid body decorated with grid-based potential maps that have been derived to reproduce results of all-atom MD simulations and refined against experimental data. Further, we have extended our grid-based formulation of interaction potentials to describe interactions between multiple SSB proteins interacting with each other, with ssDNA and with proteins represented by stationary grid potential derived from 3D cryo-electron microscopy maps. Thus, we believe our model provides a preliminary framework for accurate physical modeling of DNA replication and repair processes.

We have used several approaches to parameterize our model. Development of interaction potentials between ssDNA and SSB required substantial computational resources as it relied on brute-force sampling of ssDNA configurations around the SSB. Using enhanced sampling methods, such at the Adaptive Biasing Force [5], may considerably reduce the computational cost of the 3D potential parameterization. Our parameterization of SSB-SSB potentials, as well as the 3D potentials governing the interactions between SSB and the structurally-resolved parts of the replication fork, was done using a computationally inexpensive approach that combined an approximative treatment of van der Waals interactions and Poisson-Boltzmann electrostatics [50]. In doing so, we have neglected solvent-mediated interactions, which may be important for accurate description of the specific and nonspecific binding of biomolecules. Note that our brute-force approach accounts for the solvent mediated effects. Finally, we had to rescale our interaction potentials to reproduce the experimental values of binding free energy. We believe this step was critical for constructing a model that could quantitatively reproduce the results of force-extension experiments, something our model was not specifically parametrized to do.

While reproducing thermodynamic quantities, accurate modeling of kinetics processes remains an outstanding challenge for CG models of heterogeneous biomolecular systems, with our work being no exception. In simple terms, once a heterogeneous system has been coarse-grained, each process occurring in the system will be sped up by the smoothing of the free energy landscape. The path traveled along the free energy landscape by the system for one process might be smoothed more than for another process, which will give a larger speed up for the former. Hence, each process effectively runs on its own clock, which makes prediction or extraction of the reaction rates extremely challenging. As an example, the sliding motion of ssDNA on the SSB surface appears to be accelerated significantly more than other processes, such as the association/dissociation of ssDNA from the SSB surface. A solution to this type of problem may require extra knowledge of the kinetics of the processes of interest. In the case of SSB, one could use experimentally-known rates to slow ssDNA sliding by introducing additional friction terms that oppose DNA bead motion along the SSB surface or by adopting a stateful description of nucleotide binding the SSB surface, in either case tuning parameters to match experimental data.

For the future, we envision an extension of the model presented here, where the proteins of the replication fork are mobile and some are able to stochastically alter the state of nearby DNA, for example unwinding the DNA or synthesizing the complementary strand, in a tension-dependent manner. While such simulations may at first appear to be destined for the distant future, we believe they are within the grasp of current computational technology. Indeed, fifty years of biochemistry has provided us with extensive data regarding the individual biochemical steps, while single-molecule experiments have shown how individual biomolecules behave. Reproducing single-molecule biochemistry *in silico* will be the next enabling step toward accurate physical modeling of multi-component biological assemblies.

## Supporting information

Supplementary Animation 1

Supplementary Animation 2

Supplementary Animation 3

Supplementary Animation 4

Supplementary Animation 5

Supplementary Animation 6

Supplementary Animation 7

Supplementary Animation 8

## Acknowledgments

This work was supported by the grants from the National Science Foundation (PHY-1430124), and the National Institutes of Health (GM137015 and P41-GM104601). The supercomputer time was provided through XSEDE Allocation Grant MCA05S028 and the Leadership Resource Allocation MCB20012 on Frontera of the Texas Advanced Computing Center.

## Author Contributions

A.A. and C.M. designed the research. H-Y.C. and C.M. performed the research. C.M., H-Y.C., and A.A. wrote the manuscript.

