## Supplementary Information for "Single-molecule biophysics experiments *in silico*: Towards a physical model of a replisome"

Christopher Maffeo[a,b], Han-Yi Chou[a], Aleksei Aksimentiev[a,b]

[a] *Department of Physics, University of Illinois at Urbana–Champaign, 1110 W Green St, Urbana, 61801, IL, USA*
[b] *Beckman Institute for Advanced Science and Technology, University of Illinois at Urbana–Champaign, 405 N Matthews Ave, Urbana, 61801, IL, USA*

### Transparent Methods

#### *General simulation protocols*

All atomistic MD simulations were performed using the program NAMD [24], the TIP3P model of water [15], standard parameters for ions [3], periodic boundary conditions, particle-mesh Ewald (PME) full electrostatics with a PME grid density of about 1 Å per grid point. In the simulations of single-stranded DNA binding protein (SSB) surrounded by a solution of $dT_5$ fragments, the CHARMM36 force field [11, 6, 19, 8] was employed with NBFIX corrections applied to ion–nucleic acid interactions and to the arginine and lysine amine–nucleic acid interactions [27]. Van der Waals and short-range electrostatic energies were calculated using a smooth (7–8 Å) cutoff. Integration was performed using 2–2–6 fs multiple timestepping [2]. To enable 2-fs timestepping for bonded interactions, water bonds (and angles) and non-water covalent bonds with hydrogens were held rigid using the SETTLE [22] and RATTLE [1] algorithms, respectively.

Steric clashes that were introduced during the assembly of the system were removed through minimization using a conjugate gradient method. Subsequent simulations were performed in the NPT ensemble; a temperature of 291 K was maintained by applying Langevin forces [4] to all non-hydrogen atoms (1 $ps^{-1}$ damping coefficient) and a pressure of 1 bar was maintained by Nosé-Hoover Langevin piston pressure control [21]. The temperature was held constant using a Langevin thermostat [24] applied to all non-hydrogen atoms; the Langevin damping constant was set to 0.1 $ps^{-1}$.

All coarse-grained (CG) simulations were performed using the ARBD simulation package [5]. The two-beads-per-nucleotide model of ssDNA was implemented using the tabulated potentials, masses, damping coefficients and the all-atom mapping scheme described previously [20]. The temperature in ARBD was set to 291 K. The integration of the equations of motion was performed using and a geodesic BAOAB [18] integrator that includes a Langevin thermostat. Linear interpolation of all bonded and nonbonded potentials was employed, including three-dimensional grid-based potentials used to represent SSB proteins. The protocols of the rigid body dynamics are described in the last section of Transparent Methods.

*Generation of target data for ssDNA–SSB interaction parametrization*

As described in the main text, we performed atomic simulations to generate target distributions for the CG parametrization procedure. To enhance throughput, we five atomic models were simulated, each containing 31 $dT_5$ fragments distributed around an SSB molecule in a small cubic volume of 100 mM NaCl electrolyte, 10.6 nm on each side. The composition of the models was identical by the initial configurations were unique in each system because of the random initial arrangement of the DNA fragments. During the simulations, which lasted over 14 $\mu$s in total, the backbone atoms of residues forming $\alpha$-helices and $\beta$-sheets were harmonically restrained about their initial coordinates (1 kcal/mol Å$^2$ spring constant). The target density was obtained by mapping the all-atom representation of the DNA into a CG representation as previously described [20]. From the CG representation of the all-atom trajectories, the three-dimensional, position-dependent target density was computed for each DNA bead type and averaged over the symmetry axes of the SSB homotetramer. The target density calculations were carried out over a 2 Å grid using the volmap plugin of VMD [14] and a 4 Å-width Gaussian for each DNA bead.

*Refinement of CG SSB potentials*

An initial CG potential for each bead type was obtained by applying Boltzmann inversion to the corresponding target density, divided by a factor of ten to crudely account for the potential being applied to each bead of the same type within a CG DNA fragment. Following that, a procedure was applied to each potential to ensure that on average no force is applied at the boundary and to eliminate large forces due to extremely low bead density where the SSB core prevents access. Specifically, the potential was adjusted by uniformly subtracting the average value at the edge of the grid and then by setting the potential to 20 $k_{\mathrm{B}}T$ at voxels with a bead density $< 10^{-6}$ beads/Å$^3$. Initial trial potentials were obtained by applying Boltzmann inversion to one-tenth of the target density of that bead type.

The iterative Boltzmann inversion (IBI) [25] procedure was then applied with each iteration consisting of a CG simulation, the calculation of a CG density map, followed by an update of the

CG potential. For each iteration, CG simulations of six systems lasted a total of $\sim 80$ ns and the trajectories allowed CG density maps to be computed using the same protocol that yielded the target densities. The SSB potential for each bead type was updated as described in the main text according to the expression,

$$\Delta u = -ak_{\mathrm{B}}T \ln \frac{\rho_{\mathrm{target}} + \rho_0}{\rho_{\mathrm{CG}} + \rho_0},$$

where the scaling factor $a = 0.1$ ensured gradual convergence, and $\rho_0 = 10^{-6}$ beads/Å$^3$ protected against numerical instability. The correction to the potential was spatially smoothed using a Gaussian filter with a width of 2 Å. The potential after adding the update was adjusted using the same procedure described for the initial trial potential, ensuring no average force at the boundary and no large forces at or near the SSB core. All software for manipulating the density and potential grids was developed in-house using C++.

The density of CG DNA beads was in close agreement with all-atom MD simulations after 225 IBI iterations. The resulting potentials were used to simulate the binding of short $\mathrm{dT}_8$ fragments to SSB in CG simulations. For the purpose of our analysis, a DNA nucleotide was considered bound if its P bead was within a 4-nm-radius sphere located at the center of the SSB potential. The simulations revealed that the binding between SSB and short DNA fragments was significantly overestimated in the simulations, likely due to excess attraction in the underlying all-atom model. Hence, a procedure was enacted as described in the main text to iteratively scale the attractive regions of the P-bead potential by $\alpha_i = [1 - (f_{i-1} - 0.5)]$, where $f_{i-1}$ is the time-averaged fraction of bound nucleotides in the last iteration, until the binding fraction observed in the simulations converged to the 50% expected from experiments [17]. After scaling, the resulting potential was adjusted as described for the initial trial potentials (ensuring no average force at the boundary and no large forces at or near the SSB core).

*Motion of the SSB particle*

Upon completion of the refinement procedure, the potentials reproduced the experimentally determined binding fraction of $\mathrm{dT}_8$ fragments and, therefore, the standard binding free energy. In subsequent simulations, we modeled the SSB protein in ARBD as a rigid body having a mass and moment of inertia determined from the all-atom model. At each timestep, the force vector applied to each ssDNA bead by the grid was multiplied by $-1$ to obtain the force $\vec{F}$ applied by the bead on the SSB; the corresponding torque was obtained as a cross product of the vector connecting the center of the SSB with the DNA bead and the force vector. A Langevin force and torque [10] was added to the total force and torque to mimic the effect of the solvent with damping coefficients corresponding to the diagonal elements of the diffusion tensor obtained from Hydropro [23]. Finally, these forces and torques were used to perform integration of the rigid body's equation of motion to update position and orientation of the SSB molecule using a symplectic algorithm [7].

In the simulations that featured multiple SSB molecules, forces were evaluated between all pairs of SSB proteins using a previously described [26] protocol that clusters atoms by CHARMM36 [13] Lennard-Jones (LJ) parameters, constructs approximate 3D LJ potential maps, an electrostatic potential map from APBS [12, 16], and maps with the distribution of the corresponding LJ and electrostatic charges. If a voxel of the charge map of one SSB overlaps with the potential map of the other, a force is computed for that voxel from the product of the charge value with the negative gradient of the potential determined using a linear interpolation of the potential map; the torque is determined as a cross product between the vector leading from the origin of the SSB protein to the charge voxel and the force on that voxel. The force and the torque is scaled by $-1$ and, for the torque, transformed to apply forces and torques consistent with the Newton third law to the second SSB. Due to the relatively slow motion of the SSB proteins, the forces and torques were recalculated every 20 simulation steps.

### Supplementary Animations

**Supplementary Animation 1:** All-atom MD simulation of an SSB protein surrounded by 31 $dT_5$ ssDNA fragments. The protein (blue) and the DNA backbone (grey) are shown using a van der Waals representation, while the DNA bases (green) are shown as molecular bonds. Solvent is hidden for clarity. DNA fragments from neighboring periodic unit cell images are shown to avoid flickering associated with a periodic boundary crossing. The movie corresponds to a 250 ns MD trajectory.

**Supplementary Animation 2:** Average density of ssDNA "P" beads constructed from the results of all-atom MD simulations of the SSB protein surrounded 31 $dT_5$ DNA fragments (green semi-transparent surface) and from the results of CG simulations of the same system (purple semi-transparent surface) carried out using the CG potential maps obtained at the end of the IBI optimization procedure. The SSB core particle and the crystallographically-resolved bound

DNA (PDB accession code: 1EYG) are shown using the same representations as in Supplementary Animation 1. The animation shows a full rotation of the average density superimposed with a molecular image of the crystallographic structure about a symmetry axis of the SSB protein.

**Supplementary Animation 3:** CG simulation of a 70 nt ssDNA strand (green ball-and-stick representation) under 5 pN tension. The tension is applied in the vertical directions. The movie covers 4000 ns of the CG simulation.

**Supplementary Animation 4:** CG simulation of a 70 nt ssDNA strand (green ball-and-stick representation) under 5 pN tension with having an SSB molecule (blue molecular surface) initially bound to the DNA. The movie covers 4000 ns of the CG simulation.

**Supplementary Animation 5:** CG simulation of a 50 nt ssDNA strand (blue) attempting and failing to displace a 40 nt ssDNA strand (green) initially bound to the SSB protein (white molecular surface). The movie covers 1850 ns of the CG simulation, highlighting a failed transfer attempt.

**Supplementary Animation 6:** CG simulation of a 50 nt ssDNA strand (blue) attempting and successfully displacing a 40 nt ssDNA strand (green) initially bound to the SSB protein (white molecular surface). The movie covers 300 ns of the CG simulation, highlighting a successful transfer.

**Supplementary Animation 7:** CG simulation of the exchange of two SSB molecules (red and blue molecular surfaces) bound to the same DNA construct (green ball-and-stick representation) tethered to a surface and featuring a 70-nt ssDNA strand. The transient distortions of the dsDNA structure result from a trajectory smoothing procedure that was applied to aid visualization of the ssb transfer process. The movie covers 2100 ns of the CG simulation.

**Supplementary Animation 8:** CG simulation of the trombone loop of the T7 replication fork. Semi-transparent molecular surfaces represent the helicase (teal) in complex with the primase (purple) and polymerase (yellow) proteins, which were resolved in the cryo-EM structure of the complex [9] and represented in the CG simulation using grid-based potentials that acted on the ssDNA and SSB proteins. Double-stranded DNA and ssDNA beads bound to the above proteins are shown in the animated by were not represented explicitly in the simulations. The remaining ssDNA (green beads and rods) was coated with multiple SSB proteins (purple molecular surface). The movie covers the entire 1600 ns CG simulation.

#### Code availability

All-atom simulations were performed using the open source package NAMD, available at `https://www.ks.uiuc.edu/Research/namd/`. Coarse-grained simulations were performed using the open source package ARBD, available at `http://bionano.physics.illinois.edu/arbd`. Coarse-grained simulation configuration files for this project are available at `https://gitlab.engr.illinois.edu/tbgl/pubdata/cg-ssb`.
